# Barley (*Hordeum vulgare L*.) *HvDEP1* alleles and their effect on agronomic and physical grain traits

**DOI:** 10.64898/2026.01.27.702178

**Authors:** Hien Minh Vu, Tristan Coram, Jason Able, James Walter, Stewart Coventry, Matthew R. Tucker

## Abstract

The *Dense and erect particle 1* (*HvDEP1*) gene, located on chromosome 5H in barley (*Hordeum vulgare L*.), encodes a heterotrimeric G-protein γ-subunit that regulates grain size and stem elongation. Multiple alleles of *HvDEP1* have been identified, including the widely utilized semi-dwarf allele *HvDEP1*.*GP*, caused by an insertion mutation, and a recently discovered variant, *HvDEP1*.*V*, characterized by two deletions in the putative cis-regulatory region. In this study, we evaluated the phenotypic effects of *HvDEP1*.*V* relative to *HvDEP1*.*GP* and the wild-type allele (*HvDEP1*.*WT*) using two BC□F□ populations across multi-environment field trials spanning two locations and three years. *HvDEP1*.*V* was associated with plants that were 5-14.6 cm taller, had 3-6.7 higher lodging score, and increased head loss compared to *HvDEP1*.*GP. HvDEP1*.*V* showed comparable agronomic attributes to *HvDEP1*.*WT*. Substituting *HvDEP1*.*V* for *HvDEP1*.*GP* significantly increased all physical grain attributes, including grain width (1.44-4.24% in three out of five environments), grain length (4.88-8.69 %), grain area (6.45-11.06%) and thousand-grain weight (6.75-13.8%). Out of five environments, compared to *HvDEP1*.*WT, HvDEP1*.*V* was associated with wider grain in three environments, shorter grain in four environments, and increased grain roundness in four environments. These findings link allelic variation of the *HvDEP1* gene to key agronomic and physical grain traits and demonstrate the functional consequences of *HvDEP1*.*V* in diverse genetic backgrounds and field conditions, providing valuable insights for barley improvement.

**Key message:** By evaluating agronomic performance and physical grain traits in two genetically distinct barley populations across multiple environments, we reveal strong environment- and background-dependent effects of *HvDEP1* alleles.

## Introduction

Barley is the fourth largest crop in the world, which is used for stock feed, malt production and human food. It can be adapted to high-salinity and cold environments depending on the cultivar (Fettell et al. 2010; Munns et al. 2006; Munns and Tester 2008). Barley is also well suited to many ecogeographical zones due to the diversity of height and phenology genes, which can be manipulated to achieve adaptation. The introduction of synthetic fertilisers has led to significantly increased grain yield in crops like barley, rice and wheat. However, the stems of these crops are not strong enough to support heavy spikes, resulting in stem breakage phenomena such as lodging, brackling, head loss (Munns et al.) or necking (Berry 2019; Dockter and Hansson 2015). Since the Green Revolution, semi-dwarf genes including *Dense and erect particle 1* (*DEP1*) mutants, have been introduced into crops such as rice and wheat and have helped to increase grain yield and crop productivity (Dockter and Hansson 2015; Hedden 2003). In plants, *DEP1* encodes the gamma subunit of heteromeric G protein, which regulates stem elongation and grain size (Li et al. 2012; Sun et al. 2018). The G protein pathway consists of Gα, which provides a foundation for grain size expansion; Gβ, which is essential for plant survival and growth; and the three Gγ: *DEP1, GGC2* and *GS3* interact to regulate grain size (Sun et al. 2018). *DEP1* and *GGC2* can act on their own or together to increase grain length when bound with Gβ, while GS3 indirectly reduces grain length by competitively interacting with Gβ (Li et al. 2012).

The role of *HvDEP1* has been investigated in barley using multiple *breviaristatum* mutants in the Bowman background (Druka et al. 2011) (Table 1). The loss-of-function X-ray-induced mutation *breviaristatum-e* (*ari-e*.*GP*) or *HvDEP1*.*GP* in cv. Golden Promise shows reduced awn length, 15-20 cm shorter plant height, erect plant stature and good lodging tolerance (Dockter and Hansson 2015; Walia et al. 2007). This allele has been used extensively in European and Australian barley cultivars such as Hindmarsh, Spartacus CL and La Trobe (Jia et al. 2016). The mutation is also related to high yield and good malting quality, yet it can be associated with reduced grain size and poor weed competition (Mahajan et al. 2020; Wendt et al. 2016). Based on 12 years of yield data, *HvDEP1*.*GP* produces either more or less yield depending on the environment, demonstrating an interaction between genetic backgrounds and seasonal environments (Wendt et al. 2016).

**Table 1:**
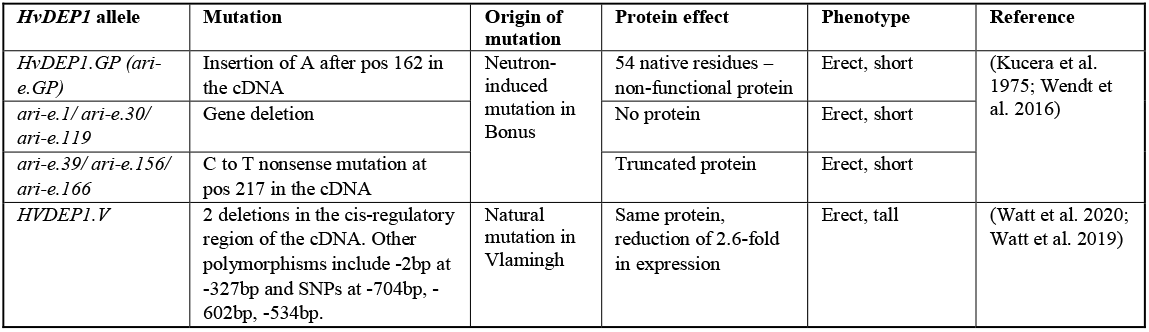
Known mutations of the *HvDEP1* gene including their origin, protein effect and phenotype.

Cultivars such as Spartacus CL and La Trobe also show good tolerance to head loss (HL), possibly as a result of *HvDEP1*.*GP* (Curry and Paynter 2019; Tucker 2022). Despite this, underlying physiological and morphological contributors to HL such as peduncle strength and length, plant height and growth habit, are poorly characterized in barley thus hindering trait improvement (Curry and Paynter 2019). Conventional phenotyping methods for HL include yield loss estimation, ground head counts, missing spike counts, and subjective rating scales. However, these methods are difficult to measure, easily confounded, labor-intensive and subject to human error. These limitations restrict breeding progress for improved HL, thus necessitating the need for new genetic targets and strategies.

A novel *HvDEP1* allele was recently identified in the Australian cultivar Vlamingh by Watt et al. (2020), containing deletions of two key cis-regulatory elements; a 9 base-pair (bp) indel (insertion-deletion mutation) and a 13 bp indel within an upstream open reading frame (Supplementary Figure S1). This results in a 2.6-fold downregulation of *HvDEP1* expression compared to the wild-type. Vlamingh was observed to be more erect than the wild-type cultivar Buloke, while showing reduced grain length and thousand-grain weight (TGW) in some environments (Watt et al. 2020; Watt et al. 2019). The agronomic and pleiotropic effects of the *HvDEP1*.*V* allele in field conditions have not been investigated or compared with the existing semi-dwarf allele, *HvDEP1*.*GP*.

In this multi-environment field study, we characterised the function of *HvDEP1* by investigating its three alleles using two genetic backgrounds: *HvDEP1*.*GP* and *HvDEP1*.*V* in an Erectoides breeding line; *HvDEP1*.*V* and *HvDEP1*.*WT* in a tall genetic background. Agronomic traits such as plant height, mechanical tolerance and yield were measured in the field. We also explored a novel phenotyping framework combining high-throughput image capture with machine learning to quantify HL in field-grown barley. Our findings indicate that the agronomic effects of *HvDEP1* are both environment- and genetic background-dependent. Furthermore, we observed that *HvDEP1*.*V* does not result in better stem breakage tolerance, while producing wider and shorter grains in some environments.

## Material and Methods

### Plant material, genotyping and multi-environment trials

This study utilised 103 genotypes including 7 relevant control cultivars, 3 parental lines and two plant populations consisting of a total of 93 BC_4_F_2_ derived lines, which were created using Vlamingh as the donor of *HvDEP1*.*V* and the recurrent parent genetic backgrounds of population 1: *HvDEP1*.*GP* from the breeding line AGTB0197; and population 2: *HvDEP1*.*WT* from the tall cultivar Beast. DNA was extracted from all genotypes and genotyped on a custom SNP chip based on an AGT proprietary linkage map. The *HvDEP1*.*V* allele was selected using an informative SNP at -704bp of *HvDEP1* at the gene’s CAF motif (Watt et al. 2020; Wendt et al. 2016). After removing the lines heterozygous for *HvDEP1*, 88 lines left can be categorised into 4 genotypic classes: (1) 28 individuals with allele *HvDEP1*.*V* and (2) 17 individuals with allele *HvDEP1*.*GP* in population 1; (3) 26 individuals with allele *HvDEP1*.*V* and (4) 17 individuals with allele *HvDEP1*.*WT* in population 2. In 2022, 103 lines were partially replicated and sown in a randomized complete block design (RCBD). To achieve full plot replication, trials in 2023 and 2024 were dropped to 72 lines, including parental lines (Beast, Vlamingh and AGTB0197), 5 relevant control cultivars (Compass, Cyclops, Flagship, RGT Planet, and Spartacus CL) and the two populations (number of lines per genotypic class are reported in Supplementary Table S2), all of which were sown in a RCBD with two replicates. The experiments were conducted with a plot size of 1.32 m × 3.2 m in two locations (Roseworthy, South Australia and Kaniva, Victoria) over three years (2022 to 2024) (Table 2). The nutrition, disease, weed and insect management were applied according to the agricultural practice of the region.

**Table 2:**
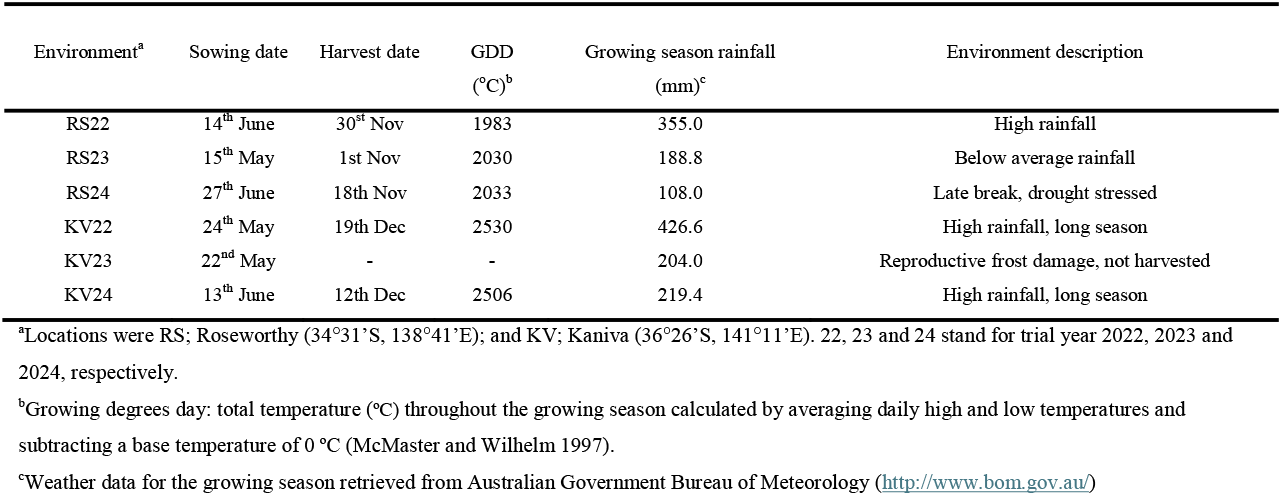
Locations and management for the multi-environment field experiments.

### Phenotyping

#### Plot-level measurements

Maturity was measured as Zadoks score (Schindelin et al.) in all environments according to the Zadoks decimal growth scale (Zadoks et al. 1974) with modifications for barley, and days to awn appearance (DAA) measured for the environments RS23 and RS24. Growth habit (GH) during stem elongation (ZS30-33) was measured using visual scores (1 = erect, 2 = semi-erect, 3 = prostrate), the degree of lodging from the base of 50% of the plot (LS) (1 = 0-10° to 9 = 80-90°) measured during grain filling, and brackling score (BS) on a 1-10 scale indicating the percentage of plot brackled. Plant height (PH) was measured as the average length of three plants within a plot from the base of the stem to the collar of the spike when spikes had fully emerged. In RS23, Normalised Difference Vegetation Index (NDVI) was collected using a GreenSeeker® 505 handheld sensor at stem elongation stage (ZS30) as a surrogate for vegetation density and crop biomass.

#### Grain characteristic measurements

Trial plots were harvested with a harvester (Quantum Plus, Wintersteiger Co., Reid, Australia) and the plot grain weight was converted to yield (kg/ha) by dividing the plot weight by the area harvested. Grain samples were cleaned over a 1.8 mm slotted sieve using a grain cleaner (Sample cleaner model SLN3, Pfeuffer GmbH, Kitzingen, Germany), and photographs processed using Fiji ImageJ (Schindelin et al. 2012) to determine the grain size attributes of grain length (GL), grain width (GW), grain area (GA), grain circularity, and grain roundness (GR). GC describes how close a shape is to a perfect circle while GR is equivalent to the inverse aspect ratio of the shape’s fitted eclipse.

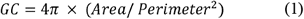

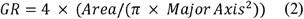

Thousand-grain weight (TGW) and hectolitre weight (HLW) were derived from the image-calculated grain number and the weight of a 30 mL subsample. Grain plumpness (GP) indicates percentage of grains retained above a 2.5 mm sieve, and screenings (SCR) represents the percentage of grains that pass through a 2.2 mm sieve.

#### Quantification of head loss using an image detection model

Head loss was quantified as counts of heads on the ground (HLC) (heads/m^2^) (Munns et al. 2006), which were counted by taking post-harvest plot images from a height of 1.3 m using Canon 70D camera with an 18 mm-lens. Prior to imaging, the plots were inspected to ensure that there was no significant loss of heads due to harvest machinery or other mechanical factors. The number of heads per image was quantified from these images using a YOLOv8 object detection model (Jocher et al. 2023), using 1,712 images (containing 44,150 total annotations) which were split into training (n = 1,396), validation (n = 256) and test sets (n = 87). The test set consisted of images which were not used in the model training process. The number of heads per m^2^ (3) was calculated by multiplying the plot area with the ground sampling distance (1), which accounts for the height of the camera at imaging, camera sensor size and field of view, then divided by the area captured per image (2).

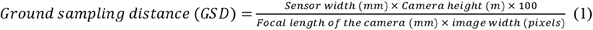

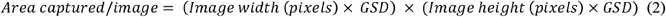

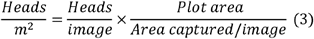

#### X-ray Computed Tomography (CT)

From the Roseworthy 2022 field trial, 10 spike samples were collected from four genotypes per genotypic class and underwent X-ray CT scanning at the Australian Plant Phenomics Facility (Adelaide, South Australia). The grain characteristics measured were the number of grains per spike, spike length, rachis internode length, grain size (mm^3^), grain weight (g), CAT radius (Wang et al. 2024) (the radius of the Circle that has the same Area as the grain maximum Transverse section) (mm), and mean grey value (average gray value of the seeds on the spike, correlated to seed density) (Hounsfield Units).

### Statistical analysis of field trials

Best linear unbiased estimates (BLUEs) of fixed effects were extracted from a mixed linear model within R/ASReml^®^ (VSN International) (Butler 2023) in the R environment, using the following general form of linear mixed model:

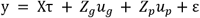

where *τ* is a vector of fixed effects with associated design matrix X; *u*_*g*_ is the vector of random Genetic by Environment effects with associated design matrix *Z*_*g*_; *u*_*p*_ is a vector of (non-genetic) random effects with associated design matrix *Z*_*p*_ and ε is the combined vector of residual errors from all environments (Smith et al. 2019). Genetic effects were modelled by explicitly fitting the gene of interest, *HvDEP1*, and genetic background as fixed variables. Non-genetic environmental variation was modelled through an AR1×AR1 correlation structure, which were fitted in the residual terms (Gilmour et al. 1997). A Tukey honest test was performed on the BLUEs using the R/biometry package (Nielsen 2025). Post-hoc Pearson correlation analysis was conducted for the Roseworthy 2023 phenotype data as this was the site with the most measured traits.

### Quantitative trait loci (QTL) mapping

To unveil the residual marker effects from the two genetic backgrounds, QTL analysis of the phenotypic traits collected from the 2022 trials was conducted on a combined analysis of both populations and on the subset of lines fixed for the *HvDEP1*.*V* allele across the different genetic backgrounds using the whole genome average interval mapping R/WGAIM (Arends et al. 2010; Verbyla et al. 2012). The package is a computational implementation of the whole genome interval mapping techniques that uses R/ASReml (Butler et al 2009) to perform statistical modeling analysis (Verbyla et al. 2007; Verbyla et al. 2012). The summary includes the magnitude of the QTL effect, the logarithm of odds (LOD) score and genetic variance (%). QTL were considered significant when LOD ≥ 3 and suggestive when LOD between 2 and 3, only QTL with a LOD greater than 2.5 were reported (He and Murabito 2014).

## Results

### Agronomic effects of HvDEP1.V

#### Population 1 (AGTB0197*4/Vlamingh)

In population 1, the *HvDEP1*.*GP* allele was associated with an erect GH (1.09–1.27), while the *HvDEP1*.*V* allele conferred a intermediate or prostrate GH (1.8–2.87) (Supplementary Table S1). *HvDEP1*.*GP* allele resulted in the shortest plant stature (34.92–83.71 cm) and the lowest lodging scores (1.3–3.5) in four out of five environments. In contrast, the *HvDEP1*.*V* allele significantly increased plant height (by 5–14.6 cm) and resulted in significantly higher lodging scores (3.0–6.7) across all environments (Fig. 1). Additionally, *HvDEP1*.*V* was associated with a yield reduction of 271–588 kg/ha compared to *HvDEP1*.*GP* in all environments.

**Fig. 1:**
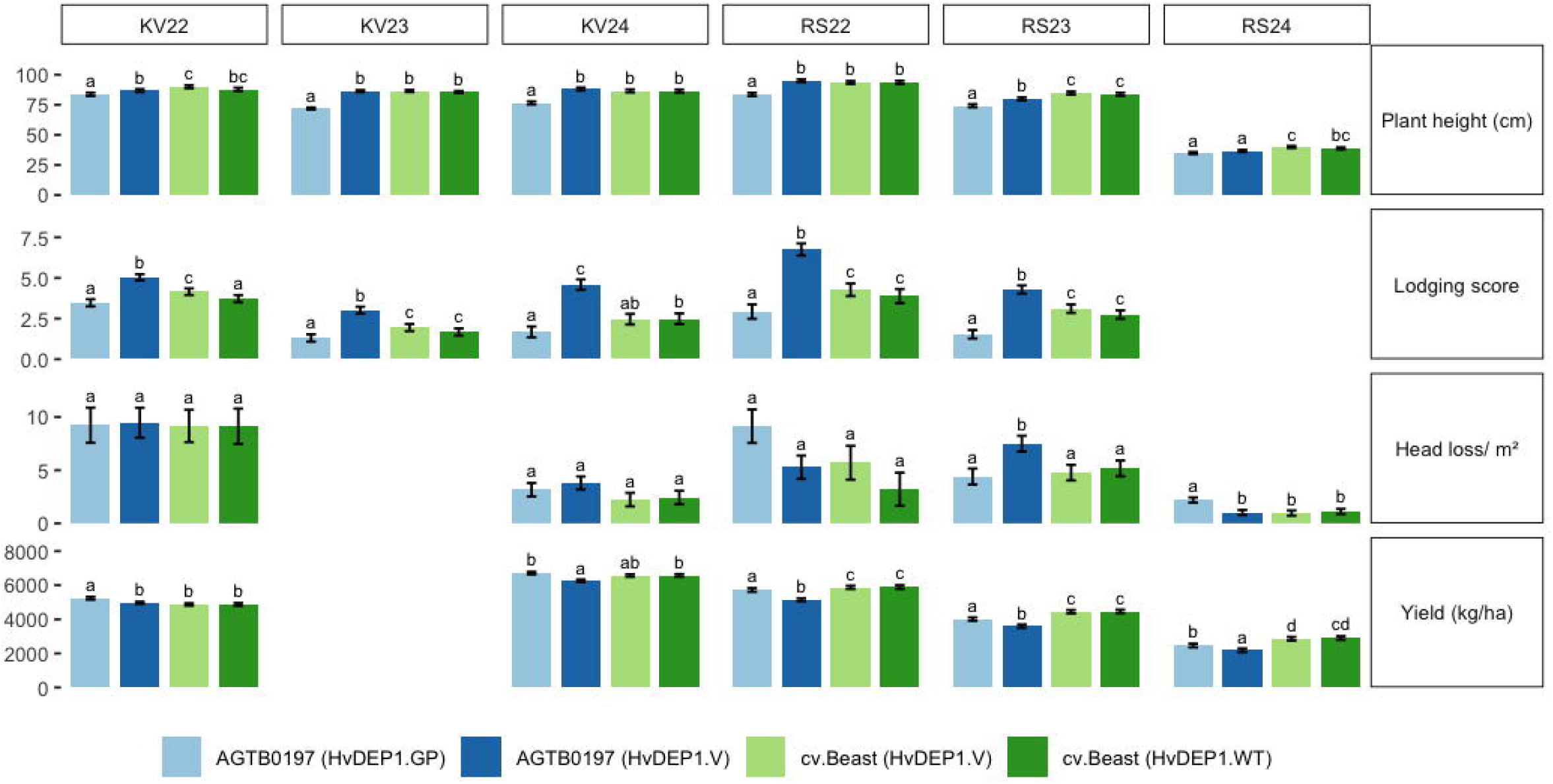
The phenotypic effect of *HvDEP1* alleles on plant height, lodging score, head loss and yield in the populations AGTB0197*4/Vlamingh and Beast*4/Vlamingh. Standard error bars are between field plot and genotype replicates with significant differences denoted by alphabetical characters.

#### Population 2 (Beast*4/Vlamingh)

During early plant development, both alleles in population 2 show a prostrate GH (2.01-2.79) (Supplementary Table S1). In most environments, when the *HvDEP1*.*WT* allele was substituted for the *HvDEP1*.*V* allele there was no significant difference in PH (up to 93.6 cm in RS22), or yield (up to 6.75 t/ha in KV24) (Fig. 1). The *HvDEP1*.*V* allele had higher LS in KV22 (4.15 compared to 3.73).

#### Head loss quantified by the image detection model

Validation metrics for the validation and test sets, predicted at a confidence of 0.30, are shown in Supplementary Figure S3a. In the test set, a correlation of 0.96 was observed between predicted and annotated heads (Supplementary Figure S3b). Head loss count (HLC) data derived from this model are reported in Fig. 1. In population 1, the *HvDEP1* allele had variable influence on HLC, with *HvDEP1*.*V* conferring the highest HLC in RS23 (7.5 heads/m^2^), and the *HvDEP1*.*GP* allele the highest HLC in RS24 (4.4 heads/m^2^), while all alleles had the same HLC in other environments. In population 2, similar HLC (up to 9 heads/m^2^ in KV22) was observed for *HvDEP1*.*V* and *HvDEP1*.*WT*.

#### HvDEP1 interactions with environment

*HvDEP1* had a significant effect on YLD, PH, and LS, but not on HLC (Table 3). Genetic background significantly influenced YLD and LS but had no detectable effect on PH or HLC. A significant interaction was observed between environment and *HvDEP1* for PH, LS, and HLC, and between environment and genetic background for YLD, PH, and LS. The effects of *HvDEP1* on YLD, PH, and LS were strongly influenced by genetic background. The only significant environment × gene × background (E×G×B) interaction was detected for LS.

**Table 3:**
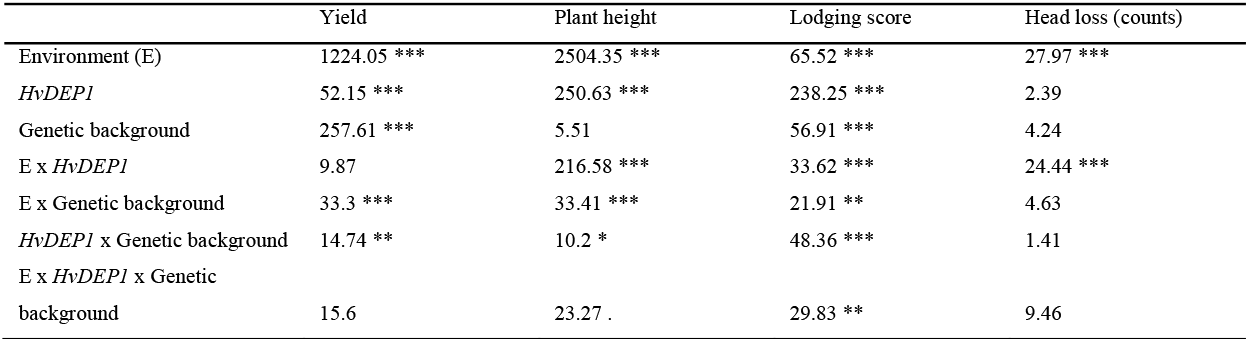
Sum of squares from multi-environment study of BC4F2 lines for 4 agronomic traits.

### Physical grain effects of HvDEP1.V

#### Population 1 (AGTB0197*4/Vlamingh)

The *HvDEP1*.*V* allele had significantly longer first and second internode length of 26.96±0.6 cm and 20.28±0.85 cm, compared to *HvDEP1*.*GP* allele with internode lengths of 20.43±0.83 cm and 17.29±1.04 cm (Fig. 2a). *HvDEP1*.*V* had significantly less grains per spike (19.87±0.59 compared to 22.28±0.61), comparable spike length, larger grain size (36.46±0.49 mm^3^ compared to 30.3±0.33 mm^3^) and heavier grain (0.057±0.001 g compared to 0.048±0.0007 g) compared to *HvDEP1*.*GP* (Fig. 2b). The *HvDEP1*.*GP* allele showed the lowest grain length, width, area and TGW compared to the other alleles (Fig. 2c & Supplementary Table S2). When *HvDEP1*.*V* was substituted for *HvDEP1*.*GP*, it significantly increased all grain attributes, 1.44-4.24% GW (3/5 environments), 4.88-8.69% GL (all environments), 6.45-11.06% GA and 6.75-13.8% TGW (all environments) (Fig. 2c, Supplementary Table S1). *HvDEP1*.*V*-carrying lines in population 1 had significantly longer grains compared to lines in population 2.

**Fig. 2.**
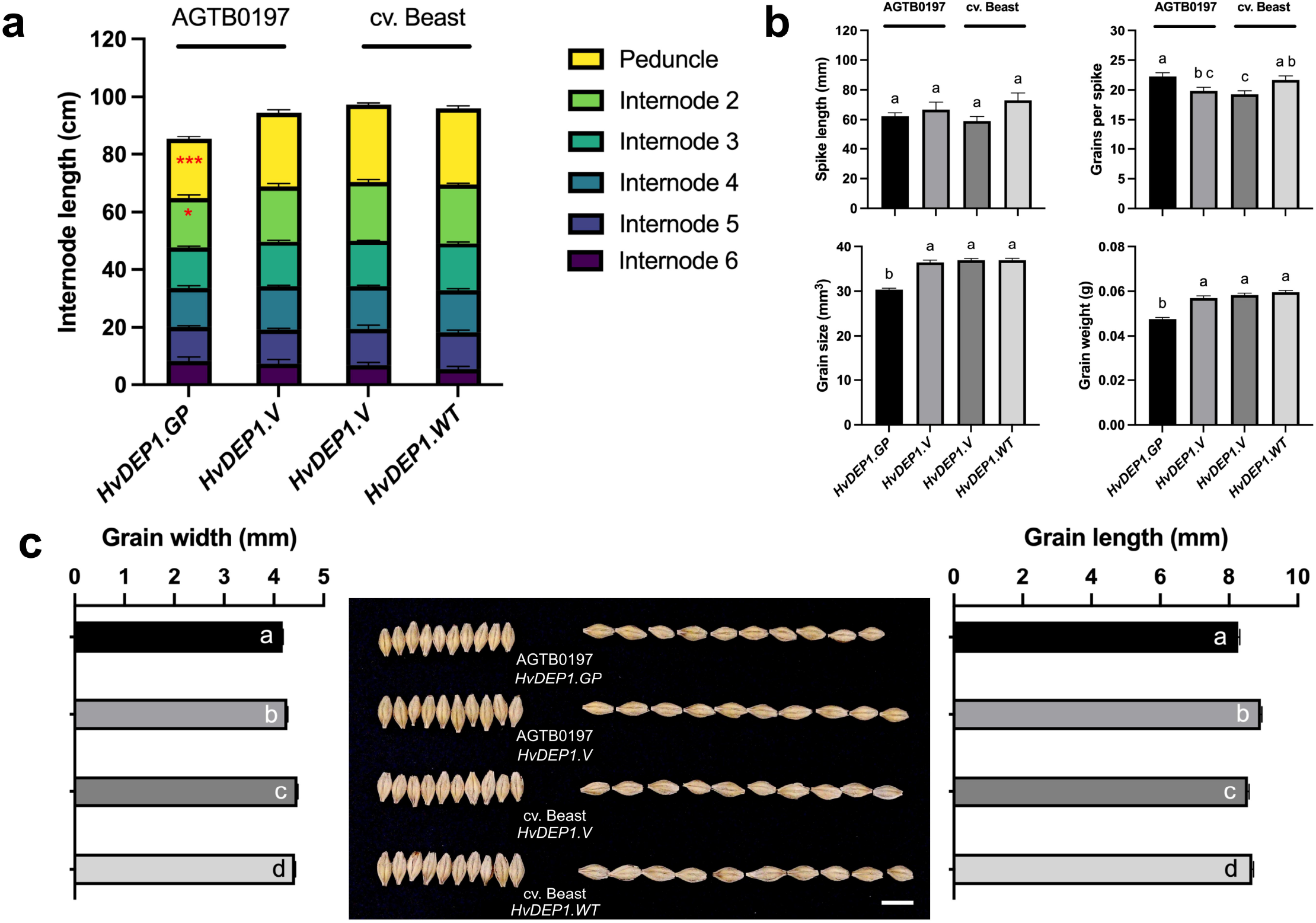
Stem, spike, and grain traits of Population 1 comparing *HvDEP1*.*GP* and *HvDEP1*.*V* alleles, and Population 2 comparing *HvDEP1*.*V* and *HvDEP1*.*WT* alleles. (a) Stem and internode lengths measured from Roseworthy 2022 samples. (b) Spike and grain traits assessed using X-ray CT from Roseworthy 2022. (c-e) Grain width and grain length from Kaniva 2022 (BLUEs predicted using ASReml-R). Scale bar represents 1 cm. Error bars indicate standard errors of the means. Significant differences are shown by distinct alphabetical groupings. Statistical significance was determined using two-way ANOVA: P < 0.05 (), P < 0.01 (), P < 0.001 (), and ns = not significant.

#### Population 2 (Beast*4/Vlamingh)

In population 2 when the *HvDEP1*.*WT* allele was substituted for *HvDEP1*.*V* there was no significant difference for internode length, ear length, grain size and grain weight (Fig. 2a & b). *HvDEP1*.*V* showed similar internode length distribution, spike length, grain size and grain weight as *HvDEP1*.*WT*. However, *HvDEP1*.*V* had fewer grains per spike (19.25±0.62 compared to 21.70±0.67) compared to *HvDEP1*.*WT* (Fig. 2b). Compared to *HvDEP1*.*WT, HvDEP1*.*V* resulted in shorter GL in RS22, RS23 and KV22; increased GW in RS22, KV22 and KV24; plumper grain in RS22 but less plump in RS23 (Fig. 2c, Supplementary Table S2).

### Genetic background effects

Based on the SNP information, on average, the progenies of population 1 (AGTB0197*4/Vlamingh) and population 2 (Beast*4/Vlamingh) are 91.57% and 88.97% genetically similar to their recurrent parent, respectively. The genetic background effects were demonstrated by identifying significant markers associated with the agronomic and grain traits (RS22 and KV22) (Supplementary Table S3) and a Pearson correlation analysis (RS23) (Fig. 3). Within the full set of lines, a total of 29 unique SNP markers were associated with 9 phenotypic traits. Notably, AGTBSNP-5H:51.03 at 51.03 cM on chromosome 5H was identified for GH, LS, YLD, TGW, GW, GA, GR and GC, representing the multi-trait effect of *HvDEP1*.*GP* allele (Supplementary Table S3.a). To elucidate the residual marker effects between the two genetic backgrounds, significant markers were identified within the *HvDEP1*.*V* subset that are different for the backgrounds. AGTBSNP-1H:128.24 (128.24 cM, LOD=2.63) on 1H, AGTBSNP-5H:48.61 (48.61 cM, LOD=2.83) on 5H and AGTBSNP-2H:73.88 (73.88 cM, LOD = 2.80) on 2H were associated for lodging score, explaining variance of 42% (RS22), 40.3% (RS22) and 46.1% (KV22) respectively. Additionally, AGTBSNP-2H:190.44 on 2H (190.44 cM, LOD=10.2, variance = 48.6%), AGTBSNP-5H:171.4 on 5H (171.4 cM, LOD=5.09, variance = 45.8%) and AGTBSNP-7H:59.22 on 7H (59.22 cM, LOD = 2.56, variance = 3.3%) explained a total of 97.7% of RS22 yield variation within this subset. For KV22 physical grain traits, AGTBSNP-3H:64.59 was identified for GR (LOD=4.33, variance = 69.3%) and GC (LOD=3.87, variance = 63.3%) in KV22, and AGTBSNP-7H:82.44 (LOD = 11.16, variance = 96%) was identified for hectolitre weight.

**Fig. 3:**
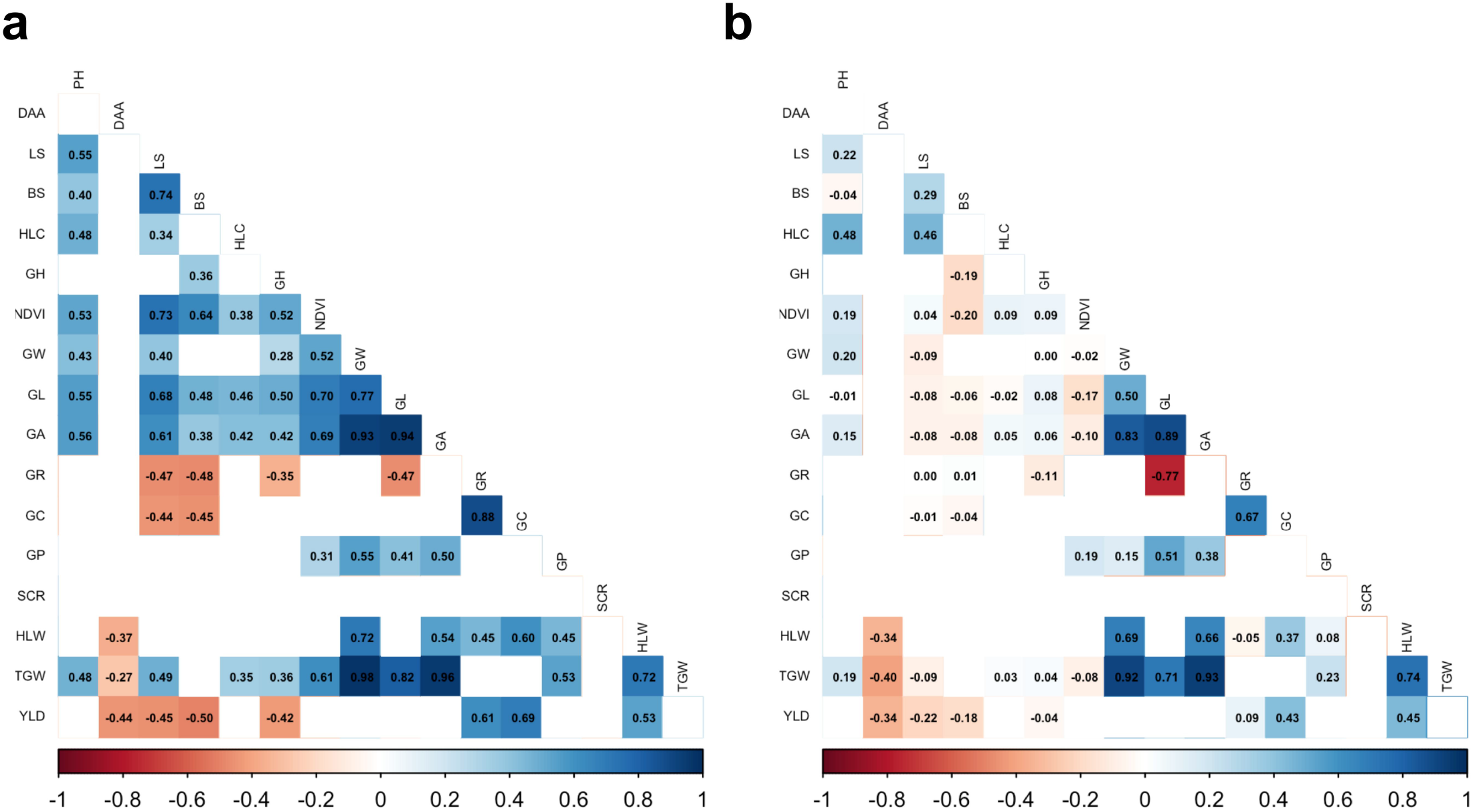
Pearson correlations of Roseworthy 2023 agronomic and grain traits within population 1 (a) AGTB0197*4/Vlamingh and population 2 (b) Beast*4/Vlamingh. Significant correlations are shown, and the non-significant correlations are left blank. PH: plant height, LS: lodging score, BS: brackling score, HLC: head loss counts, GH: growth habit score, NDVI: Normalised difference vegetation index, GW: grain width, GL: grain length, GA: grain area, GR: grain roundness, GC: grain circleness, GP: grain plumpness, SCR: screenings, HLW: Hectolitre weight, TGW: 1000-grain weight, YLD: yield.

In population 1, DAA (-0.44), LS (-0.45), BS (-0.5), GH (-0.42), and NDVI (-0.35) showed a negative correlation with yield (Fig. 3a). In contrast, GW (0.33), GR (0.61), GC (0.69), HLW (0.53), and TGW (0.25) showed a positive correlation with yield. Prostrate GH also had a positive correlation with LS and BS. Slower maturity was negatively correlated with GW, TGW, HLW and YLD. In population 2, DAA was negatively corelated with yield while GW, GL, GA, GC, HLW and TGW were positively corelated with yield (Fig. 3b). DAA was also negatively correlated with NDVI, GW, GL, GA, GC, HLW, TGW and YLD. In both populations, TGW is positively correlated with GW, GL, GA and HLW, while GP is positively associated with GL and GA.

## Discussion

### Genetic background effects

In this study, the *HvDEP1*.*V* allele from Vlamingh was backcrossed into the tall *HvDEP1*.*WT* cultivar Beast, and the erectoidies *HvDEP1*.*GP* breeding line AGTB0197, to develop two populations with contrasting *HvDEP1* alleles. Using these populations allowed comparison of the allelic effects of *HvDEP1*.*V* with *HvDEP1*.*GP* in population 1, and of *HvDEP1*.*V* with *HvDEP1*.*WT* in population 2. This backcrossing approach allowed development of populations with relatively fixed genetic backgrounds, ideal for assessing the effect of different *HvDEP1* alleles. There were potential genetic drivers from the two backgrounds, such as AGTBSNP-2H:190.44 for yield, AGTBSNP-3H:64.59 (close to *HvFT2*) for grain roundness, and AGTBSNP-7H:82.44 for hectolitre weight but further investigation is required to elucidate this effect. Though the Beast genetic background is associated with earlier maturity than AGTB0197, ranging from 1.2 to 3.2 days (Supplementary Table S1), PH and ZS did not significantly correlate (Fig. 3), and the *HvDEP1*.*V* allele has similar PH in both genetic backgrounds (Fig. 1). This indicates that within the two populations, the genetic effects on PH are independent of maturity. Both Beast and AGTB0197 contributed environment-specific QTL for some agronomic and physical grain quality traits (Supplementary Table S3). Additionally, conditions such as heat stress during grain filling period can negatively affect grain size and weight (Shirdelmoghanloo et al. 2022). Different effects of the *HvDEP1*.*V* allele, especially for GL and GW, in the different genetic backgrounds and environments, highlighted its interaction with the genetic background and sensitivity to the environmental conditions, especially during the grain filling period.

### Agronomic effects

Compared to *HvDEP1*.*GP*, the *HvDEP1*.*V* allele showed lower yield in all 5 environments, likely due to higher LS, also observed in 5 environments. In the Beast background, *HvDEP1*.*V* showed no significant difference to *HvDEP1*.*WT* in terms of yield, and no change in LS, PH, and HLC. While Vlamingh has an erect architecture, it was observed in this study that *HvDEP1*.*V* allele doesn’t contribute to the erect plant phenotype, suggesting other genetic modifiers from the Vlamingh background. Substituting *HvDEP1*.*WT* for *HvDEP1*.*V* also did not provide lodging tolerance or shorter plant height and did not have an intermediate effect when contrasted with the *HvDEP1*.*GP* allele. In terms of agronomic traits, the *HvDEP1*.*V* allele on its own has a negative effect compared to *HvDEP1*.*GP*, while it has a similar effect to *HvDEP1*.*WT*. Referring to Fig. 3, most agronomic traits negatively correlate with yield in population 1 while only DAA negatively correlates with yield in population 2. This highlights the role of plant architecture in yield improvement.

This study demonstrated that the agronomic effects of *HvDEP1* are both environment- and genetic background-dependent, particularly influencing PH, LS, HLC, and YLD. These findings underscore the importance of combining the *HvDEP1* allele with an appropriate genetic background in breeding programs. HLC varied considerably across environments and there was significant interaction between *HvDEP1* and environment for this trait, likely due to extreme conditions. For instance, RS22 and KV22 experienced high rainfall and long seasons ending with hot winds, resulting in elevated HLC across all genotypes. In contrast, RS24 was drought-stressed, leading to greater head loss in *HvDEP1*.*GP* lines compared to all lines despite having comparable height with *HvDEP1*.*V* (AGTB0197). Using an object detection model, post-harvest imagery enabled collection of HLC data. Since mechanical harvest losses can confound true HLC, pre-harvest plot inspections were conducted to ensure no significant changes occurred post-harvest. Although direct assays such as pre-harvest head counts (Curry and Paynter 2019) or wind tunnel tests (Neenan and Spencer-Smith 1975) offer more precise quantification, they are labor- and resource-intensive. The image-based method applied here provides a practical estimate of head loss under real-world mechanical harvesting conditions.

### Physical grain traits

When the *HvDEP1*.*GP* allele was substituted for *HvDEP1*.*V*, significant GL compensation was observed. Vlamingh was previously reported to have a positive grain width QTL (29.6-48.5 cM) and a negative grain length QTL (48.5-71.1 cM) on chromosome 5H, showing shorter grain in some environments (Watt et al. 2020; Watt et al. 2019). Consistent with this study, we found when the *HvDEP1*.*WT* is substituted with the *HvDEP1*.*V* allele, it results in wider grain in three environments, shorter grain in 4 environments and rounder grain in 4 environments. This indicates that the *HvDEP1*.*V* allele has an intermediate effect on grain size and shape, with the phenotype being more similar to the *HvDEP1*.*WT* allele than the *HvDEP1*.*GP* allele. Both populations showed that GL and GA positively drive important grain quality traits like TGW and GP, highlighting GL and GA as target traits for improving TGW and GP. Plumper grain and grain size uniformity are preferable in the malting industry, indicating a potential of using *HvDEP1* as a genetic tool to improve grain size and malting quality (Jiang et al. 2025; Wendt et al. 2016).

### HvDEP1.GP has pleiotropic effects on development

*HvDEP1* encodes the G protein gamma subunit and is responsible for regulating culm and seed size in barley. The gene is highly-conserved, highly expressed in meristematic tissues, has an exceptionally long 5’UTR (297bp) and is suggested to be involved in cellular growth and regulation of differentiation (Bélanger et al. 2014). Our study demonstrated that *HvDEP1* has pleiotropic effects by identifying significant marker effects at the *HvDEP1*.*GP* position, which is collocated with multiple traits like PH, LS, YLD, GA, and TGW. Similar results were observed by Wendt et al. (2016), demonstrating the collocation of DAA, PH, YLD and grain traits with *HvDEP1*.*GP*. Similarly, in rice, the *DEP1* gene has been shown to have pleiotropic effects on spike density, grain number per spike and erectness of the spike (Huang et al. 2009). The loss-of-function mutation *HvDEP1*.*GP* is associated with multi-dimensional effects due to its involvement in most meristematic tissues, but it is still unclear how *HvDEP1*.*V*, a promoter-defect allele, is involved in plant architecture modification.

In rice, the G protein pathway consists of Gα, which provides a foundation for grain size expansion; Gβ, which is essential for plant survival and growth; and the three Gγ: *DEP1, GGC2* and *GS3* interact to regulate grain size (Sun et al. 2018). *DEP1* and *GGC2* can act on their own or together to increase GL when bound with Gβ, while *GS3* indirectly reduces GL via competitively interacting with Gβ (Li et al. 2012). Additionally, *IPA1* acts as a positive regulator of *DEP1* for PH and panicle length (Jiao et al. 2010) but the two genes have contrasting effects in most yield components (Xu et al. 2014). This relationship between *DEP1* and *IPA1* offers opportunity for hybrid super rice breeding, with heterozygous plants for *DEP1* and *IPA1* showing better yield compared to homozygous plants at both loci (Xu et al. 2014). Crossing a japonica sterile line carrying desirable *DEP1* and *GS3* alleles with an indica restorer line possessing favorable *Gn1a* and *IPA1* alleles is anticipated to produce optimal yield components (Xu et al. 2016). Unravelling the G protein network in barley could provide potential for fine tuning this pathway in crop improvement.

### HvDEP1.V hints a possible uORF function

It is worth noting that the *HvDEP1*.*V* allele does not carry mutations compared to *HvDEP1*.*WT* in the coding sequence (CDS) but has deletions of two key cis-regulatory elements, possible uORFs, and is associated with a 2.6-fold lower expression of *HvDEP1* in young inflorescence tissue (Watt et al. 2020). Typically, uORFs regulate gene expression at the post-transcriptional and/or translation level, by regulating (oftentimes repressing) translation initiation of mORF and/or inducing non-sense-mediated decay in a *cis*-acting manner (Wang et al. 2024). Natural variation in the uORF of *HvDEP1*, which could be mapped as a grain length QTL, is consistent with the possibility that some QTL are uORF variants.

Notably, the *HvDEP1*.*V* allele results in fewer grains per spike (GPS) compared to both *HvDEP1*.*GP* and *HvDEP1*.*WT* (***Error! Reference source not found***.). In two-rowed barley, GPS is influenced by spikelet number, and previous studies have shown reduced GPS in HvDEP1 mutant lines such as ari-e.1 and *HvDEP1*.*GP* relative to their parental cultivars (Thirulogachandar et al. 2021; Wendt et al. 2016). *HvDEP1*.*V* also significantly alters grain traits including GL, GW, GR, and GP, with limited impact on other agronomic traits compared to *HvDEP1*.*WT*. This may reflect tissue-specific roles or expression levels of HvDEP1. Bélanger et al. (2014) reported high *HvDEP1* expression in 5 mm inflorescences but reduced levels in 20 mm stages. EoRNA data (Milne et al. 2021) show high expression in germinating embryos, root zones, and moderate expression in reproductive tissues. The OzBarley database reveals that *HvDEP1* expression in Vlamingh seedlings/germinating grain is substantially lower than in Compass, 8.91 vs. 18.85 CPM (counts per million reads), but comparable to the *HvDEP1*.*GP*-carrying Hindmarsh (9.18 CPM) (Berger et al. 2025). These findings suggest that upstream open reading frame (uORF) activity may be tissue-specific and regulated by developmental gene networks. As more transcriptomic datasets become available, integrating such resources through efficient bioinformatic pipelines will be essential for elucidating gene function.

## Conclusions

Our findings demonstrate that the *HvDEP1*.*V* allele is associated with increased plant height, reduced yield potential, and decreased stem integrity compared to the widely adopted *HvDEP1*.*GP* semi-dwarf allele. Despite these drawbacks, *HvDEP1*.*V* conferred significant increases in grain width, length, area, and thousand-grain weight in several environments, collaborating the results of the previous study by Watt et al. (2020). Relative to the wild-type allele (*HvDEP1*.*WT*), *HvDEP1*.*V* exhibited similar phenotypic profiles, though it was associated with fewer grains per spike, increased head loss counts, wider grain, and shorter grain in specific environments.

Future research should investigate the interaction of *HvDEP1*.*V* with other semi-dwarfing genes and its performance across diverse genetic backgrounds to further elucidate its breeding potential. Incorporation of *HvDEP1*.*V* into breeding programs may be feasible when combined with erect plant architecture or complementary dwarfing loci. Additionally, detailed analyses of the regulatory modifications and tissue-specific expression patterns of *HvDEP1*.*V* will help clarify the gene’s functional mechanisms and support more targeted deployment in barley improvement.

## Acknowledgements

We acknowledge the Adelaide University Stipend for the PhD scholarship and Australian Plant Breeding Academy for the supplementary scholarship. We thank Australian Grain Technologies for their contribution in population development, genotyping and field experiments management. We also acknowledge the use of the facilities, and scientific and technical assistance of the Australian Plant Phenomics Facility, which is supported by the Australian Government’s National Collaborative Research Infrastructure Strategy (NCRIS).

## Statements & Declarations

## Funding

The authors declare that no funds, grants, or other support were received during the preparation of this manuscript.

## Conflicts of interest/Competing interests

On behalf of all authors, the corresponding author states that there is no conflict of interest.

## Author contribution statement

HMV contributed to the experimental process, data analysis, results interpretation and manuscript preparation. SC conceptualised the study, prepared the plant materials and coordinated the experimental process. JW developed the head loss object detection model described in the method. All authors were involved in analysis interpretation, manuscript editing and reviewing. All authors read and approved the final manuscript.

## Data availability statement

The phenotype datasets generated and analysed during the current study are available from the corresponding author on reasonable request.

## Supplemental data

1. Supplementary Table S1: Multi-environment BLUEs of agronomic traits associated with two genetic backgrounds and three HvDEP1 alleles. Agronomic traits collected in field trials including maturity, yield, plant height, lodging score, brackling score, head loss. Significant differences are annotated into groups as different alphabetical letters using post-hoc Tukey test
2. Supplementary Table S2: Multi-environment BLUEs of grain traits associated with population genetic background and HvDEP1 alleles. Significant differences are annotated into groups as different alphabetical letters using post-hoc Tukey test
3. Supplementary Table S3: QTL associated with agronomic and grain characteristics in two biparental populations across multiple environments
4. Supplementary Figure S1: Genetic structure of HvDEP1.GP and HvDEP1.V alleles compared to the wildtype, adapted from Watt et al. (2020)
5. Supplementary Figure S2: Spikes and grain traits from X-ray CT
6. Supplementary Figure S3: Head loss detection model (A) Accuracy graph of the images predicted (B) Validation metrics of the head loss detection model trained in Yolov8 for the validation and test sets

**Supplementary Figure S1.**
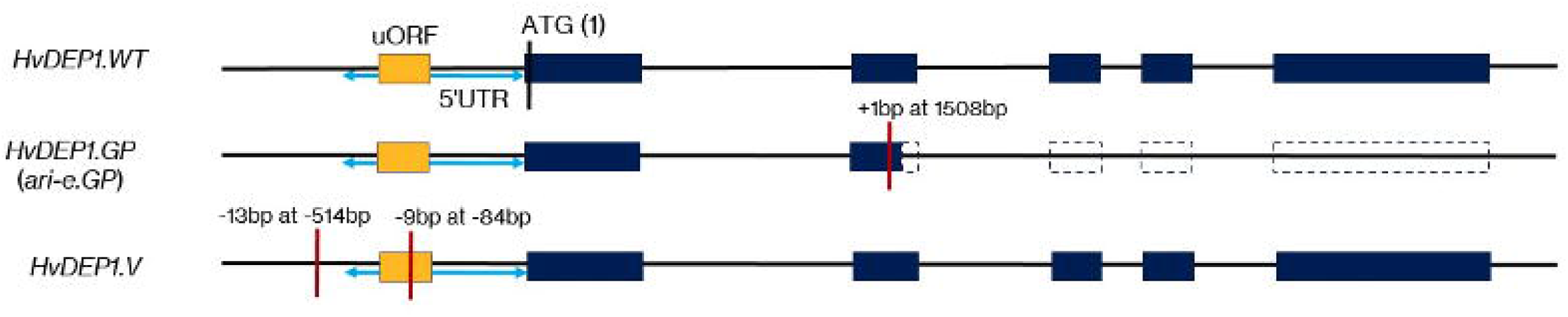
Genetic structure of HvDEP1.GP and HvDEP1.V alleles compared to the wildtype, adapted from Watt et al. (2020)

**Supplementary Figure S2.**
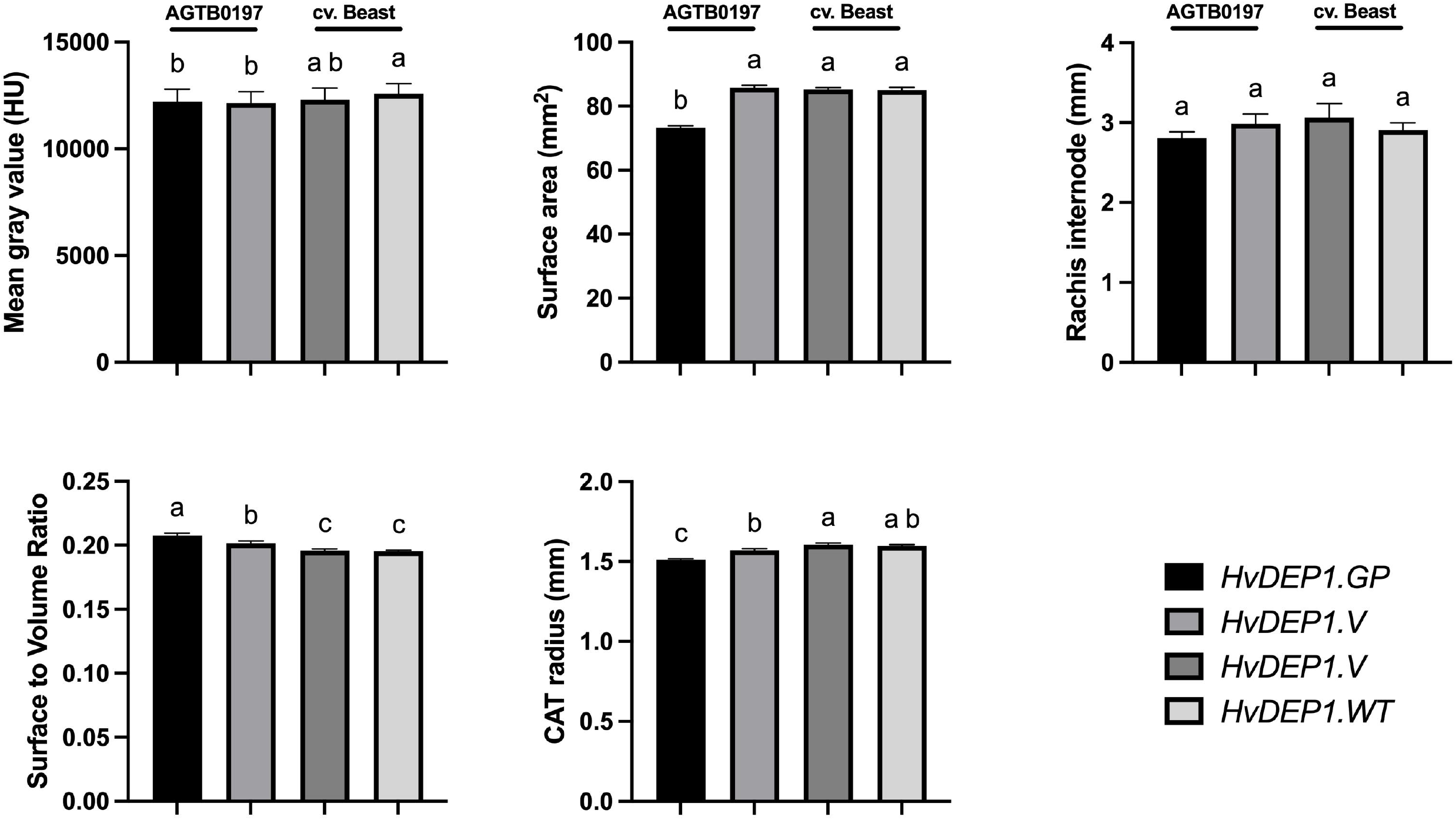
Spikes and grain traits from X-ray CT. Mean gray value (Hounsfield unit) = average gray value of the seeds on the spike, correlates to seed density; CAT radius = radius of the circle that has the same area as the seed maximum transverse section. Error bars indicate standard error between 5 genotype replicates x 3 technical replicates. Significant differences are denoted as alphabetical characters, as determined by one-way ANNOVA test

**Supplementary Figure S3.**
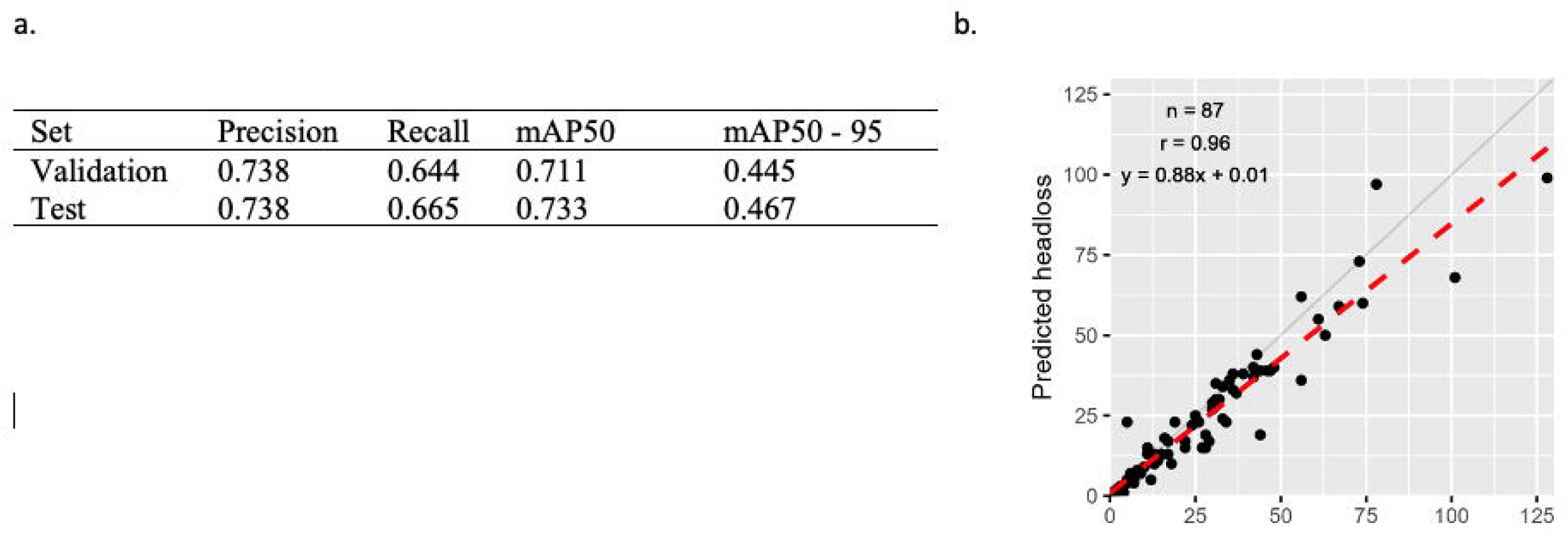
Head loss detection model (a) Accuracy graph of the images predicted (b) Validation metrics of the head loss detection model trained in Yolov8 for the validation and test sets

## Notes

### Competing Interest Statement

The authors have declared no competing interest.

## References

Arends D, Prins P, Jansen RC, Broman KW (2010) R/qtl: high-throughput multiple QTL mapping. Bioinformatics 26:2990–2992. 10.1093/bioinformatics/btq565

Bélanger S, Gauthier M, Jean M, Sato K, Belzile F (2014) Genomic characterization of the Hordeum vulgare DEP1 (HvDEP1) gene and its diversity in a collection of barley accessions. Euphytica 198:29–41.10.1007/s10681-014-1089-1

Berger B, Tucker M, Baumann U, Kalashyan E, Schwerdt J, Chalmers K (2025) OzBarley: genotypic and expression data of the OzBarley elite panel. In: Adelaide TUo (ed)10.25909/29104916.v1

Berry PM (2019) Lodging Resistance in Cereals. In: Savin R, Slafer GA (eds) Crop Science. Springer New York, New York, NY, pp 209–227. 10.1007/978-1-4939-8621-7_228

Butler DG, Cullis, B.R., Gilmour, A.R., Gogel, B.G., Thompson, R (2023) ASReml-R Reference Manual Version 4.2. VSN International Ltd., Hemel Hempstead, HP2 4TP, UK.

Curry J, Paynter B (2019) Causes and impact of barley head loss in the Western Region, GRDC.10.13140/RG.2.2.11179.05921

Dockter C, Hansson M (2015) Improving barley culm robustness for secured crop yield in a changing climate. J Exp Bot 66:3499–3509. 10.1093/jxb/eru521

Druka A, Franckowiak J, Lundqvist U, Bonar N, Alexander J, Houston K, Radovic S, Shahinnia F, Vendramin V, Morgante M, Stein N, Waugh R (2011) Genetic dissection of barley morphology and development. Plant Physiol 155:617–627. 10.1104/pp.110.166249

Fettell N, Bowden P, McNee T, Border N (2010) Barley growth & development. Industry & Investment NSW

Gilmour AR, Cullis BR, Verbyla AnP (1997) Accounting for Natural and Extraneous Variation in the Analysis of Field Experiments. Journal of Agricultural, Biological, and Environmental Statistics 2:269–293.10.2307/1400446

He C, Murabito JM (2014) Genome-wide association studies of age at menarche and age at natural menopause. Mol Cell Endocrinol 382:767–779. 10.1016/j.mce.2012.05.003

Hedden P (2003) The genes of the Green Revolution. Trends Genet 19:5–9. 10.1016/s0168-9525(02)00009-4

Huang X, Qian Q, Liu Z, Sun H, He S, Luo D, Xia G, Chu C, Li J, Fu X (2009) Natural variation at the DEP1 locus enhances grain yield in rice. Nat Genet 41:494–497. 10.1038/ng.352

Jia Q, Tan C, Wang J, Zhang XQ, Zhu J, Luo H, Yang J, Westcott S, Broughton S, Moody D, Li C (2016) Marker development using SLAF-seq and whole-genome shotgun strategy to fine-map the semi-dwarf gene ari-e in barley. BMC Genomics 17:911. 10.1186/s12864-016-3247-4

Jiang C, Kan J, Gao G, Dockter C, Li C, Wu W, Yang P, Stein N (2025) Barley2035: A decadal vision for barley research and breeding. Mol Plant 18:195–218. 10.1016/j.molp.2024.12.009

Jiao Y, Wang Y, Xue D, Wang J, Yan M, Liu G, Dong G, Zeng D, Lu Z, Zhu X, Qian Q, Li J (2010) Regulation of OsSPL14 by OsmiR156 defines ideal plant architecture in rice. Nat Genet 42:541–544.10.1038/ng.591

Jocher G, Chaurasia A, Qiu J (2023) Ultralytics YOLOv8.

Kucera J, Lundqvist U, Gustafsson A (1975) Induction of breviaristatum mutants in barley. Hereditas 80:263–278. 10.1111/j.1601-5223.1975.tb01525.x

Li S, Liu Y, Zheng L, Chen L, Li N, Corke F, Lu Y, Fu X, Zhu Z, Bevan MW, Li Y (2012) The plant-specific G protein gamma subunit AGG3 influences organ size and shape in Arabidopsis thaliana. New Phytol 194:690–703. 10.1111/j.1469-8137.2012.04083.x

Mahajan G, Hickey L, Chauhan BS (2020) Response of Barley Genotypes to Weed Interference in Australia. Agronomy 10:99. 10.3390/agronomy10010099

McMaster GS, Wilhelm WW (1997) Growing degree-days: one equation, two interpretations. Agricultural and Forest Meteorology 87:291–300. 10.1016/S0168-1923(97)00027-0

Milne L, Bayer M, Rapazote-Flores P, Mayer C-D, Waugh R, Simpson CG (2021) EORNA, a barley gene and transcript abundance database. Scientific Data 8:90. 10.1038/s41597-021-00872-4

Munns R, James RA, Lauchli A (2006) Approaches to increasing the salt tolerance of wheat and other cereals. J Exp Bot 57:1025–1043. 10.1093/jxb/erj100

Munns R, Tester M (2008) Mechanisms of salinity tolerance. Annu Rev Plant Biol 59:651–681.

Neenan M, Spencer-Smith JL (1975) An analysis of the problem of lodging with particular reference to wheat and barley. The Journal of Agricultural Science 85:495–507. 10.1017/S0021859600062377

Nielsen SR, Sam; Conway, Annie (2025) biometryassist: Functions to Assist Design and Analysis of Agronomic 76 Experiments.

Schindelin J, Arganda-Carreras I, Frise E, Kaynig V, Longair M, Pietzsch T, Preibisch S, Rueden C, Saalfeld S, Schmid B, Tinevez J-Y, White DJ, Hartenstein V, Eliceiri K, Tomancak P, Cardona A (2012) Fiji: an open-source platform for biological-image analysis. Nature Methods 9:676–682. 10.1038/nmeth.2019

Shirdelmoghanloo H, Chen K, Paynter BH, Angessa TT, Westcott S, Khan HA, Hill CB, Li C (2022) Grain-Filling Rate Improves Physical Grain Quality in Barley Under Heat Stress Conditions During the Grain-Filling Period. Front Plant Sci 13:858652. 10.3389/fpls.2022.858652

Smith AB, Borg LM, Gogel BJ, Cullis BR (2019) Estimation of Factor Analytic Mixed Models for the Analysis of Multi-treatment Multi-environment Trial Data. Journal of Agricultural, Biological and Environmental Statistics 24:573–588. 10.1007/s13253-019-00362-6

Sun S, Wang L, Mao H, Shao L, Li X, Xiao J, Ouyang Y, Zhang Q (2018) A G-protein pathway determines grain size in rice. Nat Commun 9:851. 10.1038/s41467-018-03141-y

Thirulogachandar V, Koppolu R, Schnurbusch T (2021) Strategies of grain number determination differentiate barley row types. Journal of Experimental Botany 72:7754–7768. 10.1093/jxb/erab395

Tucker MR (2022) Revealing the basis for head loss in barley, SAGIT.

Verbyla AP, Cullis BR, Thompson R (2007) The analysis of QTL by simultaneous use of the full linkage map. Theor Appl Genet 116:95–111. 10.1007/s00122-007-0650-x

Verbyla AP, Taylor JD, Verbyla KL (2012) RWGAIM: an efficient high-dimensional random whole genome average (QTL) interval mapping approach. Genet Res (Camb) 94:291–306. 10.1017/S0016672312000493

Walia H, Wilson C, Condamine P, Ismail AM, Xu J, Cui X, Close TJ (2007) Array-based genotyping and expression analysis of barley cv. Maythorpe and Golden Promise. BMC Genomics 8:87. 10.1186/1471-2164-8-87

Wang J, Liu J, Guo Z (2024) Natural uORF variation in plants. Trends in Plant Science 29:290–302. 10.1016/j.tplants.2023.07.005

Watt C, Zhou G, Angessa TT, Moody D, Li C (2020) A novel polymorphism in the 5′ UTR of HvDEP1 is associated with grain length and 1000-grain weight in barley (Hordeum vulgare). Crop and Pasture Science 71 10.1071/cp20169

Watt C, Zhou G, McFawn LA, Chalmers KJ, Li C (2019) Fine mapping of qGL5H, a major grain length locus in barley (Hordeum vulgare L.). Theor Appl Genet 132:883–893. 10.1007/s00122-018-3243-y

Wendt T, Holme I, Dockter C, Preuss A, Thomas W, Druka A, Waugh R, Hansson M, Braumann I (2016) HvDep1 Is a Positive Regulator of Culm Elongation and Grain Size in Barley and Impacts Yield in an Environment Dependent Manner. PLoS One 11:e0168924. 10.1371/journal.pone.0168924

Xu H, Zhao M, Zhang Q, Xu Z, Xu Q (2016) The DENSE AND ERECT PANICLE 1 (DEP1) gene offering the potential in the breeding of high-yielding rice. Breed Sci 66:659–667. 10.1270/jsbbs.16120

Xu Q, Xu N, Xu H, Tang L, Liu J, Sun J, Wang J (2014) Breeding value estimation of the application of IPA1 and DEP1 to improvement of Oryza sativa L. ssp. japonica in early hybrid generations. Molecular Breeding 34:1933–1942. 10.1007/s11032-014-0150-z

